# Myelin water fraction decrease in mild traumatic brain injury

**DOI:** 10.1101/2020.02.05.934430

**Authors:** Bretta Russell-Schulz, Irene Vavasour, Jing Zhang, Alex L. MacKay, Victoria Purcell, Angela M. Muller, Leyla Brucar, Ivan J. Torres, William Panenka, Naznin Virji-Babul

## Abstract

The increased incidence of reported traumatic brain injury (TBI) and its potentially serious long-term consequences have enormous clinical and societal impacts. The diffuse and continually evolving secondary changes after TBI make it challenging to evaluate the changes in brain-behaviour relationships. In this study we used myelin water imaging to evaluate changes in myelin water fraction (MWF) in individuals with chronic brain injury and evaluated their cognitive status using the NIH Toolbox Cognitive Battery. Twenty-two adults with mild or severe brain injury and twelve age, gender and education matched healthy controls took part in this study. We found a significant decrease in global white matter MWF in individuals with mild TBI compared to the healthy controls. Significantly lower MWF was evident in most white matter ROIs examined including the corpus callosum (separated into genu, body and splenium), minor forceps, right anterior thalamic radiation, left inferior longitudinal fasciculus; and right and left superior longitudinal fasciculus and corticospinal tract. No significant correlations were found between MWF in mild TBI and the cognitive measures. These results show for the first time the loss of myelin in chronic mild TBI.

## 1 Introduction

Every year about 160,000 Canadians sustain brain injuries, the most prevalent causes being falls and motor vehicle accidents (MVA) (Brain Injury Canada). An impact to the head results in an immediate and direct insult to the brain, setting off a complex cascade of metabolic and neurochemical events. These effects can lead to long-term changes in brain physiology leading to cognitive, motor and affective dysfunction (1–3). Over a lifetime, repeated brain trauma or a single moderate or severe TBI is associated with an increased incidence of multiple neuropsychiatric conditions and is a significant risk factor for developing neurodegenerative disorders (4,5). The diffuse and continually evolving secondary changes that are the hallmark of TBI have made it extremely challenging to evaluate brain-behavior relationships. Neurodegeneration has been found in the chronic phases after TBI(6) and detected using atrophy measures, however conventional neuroimaging tools (such as CT and MRI) cannot detect the widespread and often subtle changes in structure and function that occur in the brain following TBI (7). The long-term effects of a mild TBI (mTBI) are less clear than more severe head injuries and neuroimaging findings are less consistent(8,9). Thus advanced MR imaging techniques are increasingly being used to examine and monitor changes in the brain following TBI (10–12).

Physiologically, movement of the brain within the skull causes shearing/stretching of the axons and initiates a cascade of molecular events which disrupts normal brain cell function. Metabolic changes occur rapidly following axonal strain, altering the permeability of sodium channels, resulting in an increase in intra-axonal calcium and ultimately a failure of the sodium pump, causing further metabolic disruptions (13). Long white matter tracts are particularly vulnerable to these forces and diffuse axonal injury (DAI) can lead to demyelination and axonal loss resulting in brain atrophy (5,14–18). A long term follow-up study of individuals with severe TBI found neurodegeneration in the chronic phase as measured by volume reduction in the corpus callosum between 1-8 years post injury (19). Loss of myelin increases axonal vulnerability to further trauma and may predispose the axon to further damage (20,21). In animal models of TBI, extensive demyelination is consistently reported (22,23), and human post mortem studies have found evidence of loss of myelin (24) years after the initial TBI symptoms have resolved. Given the importance of monitoring white matter tracts in TBI, a validated quantitative technique to examine myelination (loss and possible remyelination (20,25)) of these tracts would provide a powerful approach to understand the pathology of TBI, the evolving nature of the injury and ultimately could be used to evaluate the impact of neurorehabilitation.

Measuring myelin *in* vivo using MR is nontrivial and requires specialized techniques (26,27). Myelin is a lipid rich membrane assembly that wraps around axons in a series of layers. Myelin enables saltatory conduction which produces fast and efficient neuronal signal transmission across the brain and throughout the body. The loss or disruption of myelin results in loss of saltatory conduction causing slower signal propagation and thus delay of translation of information. Measuring non-aqueous protons in myelin is difficult because the MR signal decays too rapidly to be measured by common MR techniques. The water trapped between myelin bilayers is known as myelin water and it is accessible using MR. Several different techniques have been developed to measure the myelin water (28), one such approach involves using multi-component T2 analysis, myelin water imaging (MWI). Multi-component T2 relaxation can be used to detect different water environments in human brain tissue (29–31), including a short T2 component that is thought to be myelin water (32–40). The proportion of water in brain that can be attributed to myelin water is known as the myelin water fraction (MWF) which can be used as a marker for myelin (41). MWF has been validated as a myelin marker in histological studies in animal models (42–44) and human post-mortem multiple sclerosis brain (45–47) and spinal cord (48). MWF has been found to be decreased in normal appearing white matter in neurodegenerative diseases such as multiple sclerosis (49–52). MWF has previously been shown to decrease post single sports concussion (mTBI) and then recover after 2 months (53), showing the utility of MWF in mTBI.

It should be noted that there are other factors aside from pathology that can cause changes to myelination and thus MWF. In humans, myelin development begins in utero and develops quickly through the first few years but continues to increase through adolescence and young adulthood (54) and is thought to increase with learning in healthy adult brains (55). MWF in healthy individuals is significantly correlated with increasing age and years of education (56,57). A subsequent study (58) confirmed these significant correlations with healthy control MWF but also found age and education relationships in a cohort of first-episode schizophrenia patients.

Cognitive decline is a common consequence of moderate/severe TBI and can continue long-term (59). In most cases of mTBI, there is little to no evidence of long-term cognitive impairment(60–63), however a portion of patients with mTBI will report persistent symptoms, such as post-concussive syndrome and other cognitive function issues(64–69). It is important to characterize the cognitive function of TBI subjects as it is a potent predictor of functional capacity, and targeted rehabilitation measures are available. A number of groups have documented an association between myelin water fraction and cognition. Lang *et al.*(58) showed that Frontal white matter MWF correlated with scores on a premorbid IQ test, the North American Adult Reading Test (NAART) and Choi *et. al.* (70) found a correlation between a processing speed index and a global measure of apparent myelin water fraction in a group of moderate to severe TBI subjects 3 months post-injury. These findings show that myelin measures may reflect cognitive functional changes in TBI.

Here we examine myelin water fraction (MWF) within a heterogeneous group of chronic TBI participants and compare mild TBI MWF to healthy controls using a whole cerebrum coverage MWF sequence to characterize differences in myelin. We also examined the effects of age and years of education on MWF, independent of TBI. Finally, we compared cognitive scores between the two groups and their relationship with MWF. Based on results from a previous study in TBI (70) we hypothesized that MWF would be particularly associated with fluid cognitive abilities.

## 2 Methods

### 2.1 Subject demographics

Participants were recruited from brain injury associations across the Greater Vancouver area as part of a study to evaluate the impact of an intensive cognitive intervention program (Arrowsmith) in adults with TBI (71). All healthy controls were recruited from the lower mainland of Vancouver in close proximity to the university. All controls were screened to ensure that they had no history of head trauma, neuropsychiatric disorders, or any other neurological conditions. The participants with TBI were first interviewed to determine eligibility and to evaluate the severity of TBI. Participants were excluded if they were currently in litigation or if they had other severe medical conditions affecting brain function. They were also excluded if they had a diagnosis of psychiatric illness based on a Mini-international neuropsychiatric interview (MINI) (72). The severity of injury was based on self-report using the diagnostic criteria from the World Health Organization (WHO). Information was retrospectively reported by the subjects, where mild TBI was defined as Loss of Consciousness (LOC) of less than 30 minutes and Post-traumatic amnesia (PTA) less than 24 hours, and severe TBI was defined as LOC over 24 hours. All participants provided written consent according to the guidelines set forth by the Clinical Research Ethics Board at the University of British Columbia. Subject demographics are given in Table 1, and individual subject TBI history and severity are given in Supplementary Table S1. Time since injury varied from 6 months to 28 years post injury across subjects, and 6 months was assumed to be in the chronic stage for mTBI(73).

**Table 1:** Subject demographics; where TBI = Traumatic Brain Injury and m=mild and s=severe. Age and education compared between controls and mTBI using an independent samples t-test and a chi-squared test to compare group gender ratios

| Subject Group | N | Gender Ratio (M:F) |  | Age (years) Mean±sd (Range) |  | Education (years) Mean±sd (Range) |  |
| --- | --- | --- | --- | --- | --- | --- | --- |
| Control | 12 | 8:4 | p=0.8 | 37.2±12 (21-53) | p=0.8 | 16.4±3 (12-20) | p=0.2 |
| mTBI | 18 | 9:9 |  | 38.5±12 (18-57) |  | 15.5±2 (12-18) |  |
| sTBI | 4 | 3:1 |  | 47.5±6 (40-54) |  | 14.5±2 (12-18) |  |

### 2.2 MRI protocol

MRI scans were completed on a 3T Philips Achieva scanner. Sequences included a 3DT1 structural scan (MPRAGE TR=3000 ms, TI = 1072 ms, 1×1×1mm^3^ voxel, 160 slices) for registration and segmentation of global white matter and white matter regions of interest (ROIs) and a 3D 48-echo Gradient and Spin Echo (GRASE) T2 relaxation sequence with an EPI factor of 3 (TR=1073, echo spacing = 8ms, 20 slices acquired at 1 × 2 × 5 mm and 40 slices reconstructed at 1 × 1 × 2.5 mm, FOV 230 × 190 × 100 mm, acquisition time 7.5 min) for MWF determination (74,75).

### 2.3 Neuropsychological testing

The NIH Toolbox Cognition Battery (76) was used to evaluate cognition in both groups. The Test of Memory Malingering was used to exclude individuals showing suboptimal effort. Two primary composite cognitive measures were employed in this study (77).The Crystallized composite score (age adjusted) from the NIH Toolbox was used to assess crystallized cognitive ability, which refers to an accumulated storage of verbal knowledge and skills that are heavily influenced by educational and cultural experience. The crystallized score was based on the average performance on the Picture Vocabulary Test (language) and Oral Reading Recognition Test (language). The Fluid composite score was based on performance on the following NIH Toolbox measures: Dimensional Change Card Sort Test (attention, executive function), Flanker Inhibitory Control and Attention Test (attention, executive function), Picture Sequence Memory Test (episodic memory), Pattern Comparison Processing Speed Test (processing speed), List Sorting Working Memory Test (working memory). Fluid ability is defined as the capacity for new learning and information processing in novel situations, which is especially influenced by biological processes and is less dependent on past exposure. All scores were age-adjusted based on the NIH Toolbox nationally representative U.S. normative sample, and standard scores had a mean of 100 and standard deviation of 15. Three healthy controls did not complete cognitive testing, one due to familiarity with the tests.

### 2.4 Myelin water fraction (MWF) data analysis

The signal decay curve obtained by the T2 relaxation sequence was modelled by multiple exponential components and the T2 distribution was estimated using in-house software in matlab (MathWorks, Massachusetts, U.S.A) which contained a regularized non-negative least squares algorithm using the extended phase graph and flip angle estimation to deal with stimulated echo artifacts (29–31,78,79). MWF in each image voxel was computed as the ratio of the area under the T2 distribution with times of 10-40ms to the total area under the distribution. MWF and 3DT1 images were registered to MNI space and brain extracted and segmented using tools from FSL (80–82). Slices below the inferior portion of the hypothalamus (mammillary body) were rejected in order to account for differing head sizes and make sure MWF was measured over the same brain coverage for all subjects. Global white matter (GWM) was segmented and pre-existing regions of interest from the JHU tract atlas in FSL (83–85) were used to segment specific tracts using FSL: splenium of corpus callosum (SCC), body of corpus callosum (BCC), genu of corpus callosum (GCC), minor forceps (MN), and right and left anterior thalamic radiation (ATR), inferior and superior longitudinal fasciculus (ILF and SLF respectively), and corticospinal tract (CST).

### 2.5 Statistical analysis

All statistical analysis was completed using IBM SPSS Statistics Version 25. Demographic differences were examined; age and education were compared between groups using an independent samples t-test and a chi-squared test was used to compare gender ratios between groups, where p<0.05 was considered significant. All group comparisons were done between controls and mild TBI, due to the low subject numbers in the severe TBI category. Mean MWF for ROIs were compared between participant groups using an independent samples t-test, where p<0.05 was considered significant. Bonferroni-Holm correction for multiple comparisons was used to correct for multiple comparisons. The association between age and years of education and MWF in each ROI was examined using Pearson’s correlations across controls and mTBI. A linear model comparing MWF (ANCOVA) across groups was used with age and education as covariates. Finally, cognitive scores were compared between groups using an independent sample’s t-test and a Cohen’s d to assess effect size. MWF was compared with cognitive scores in mTBI using Pearson’s correlation.

## 3 Results

### 3.1 Demographic characteristics between groups

Table 1 shows the general group demographics and Supplementary Table S1 gives demographic and clinical features for all the participants in this study. Age, education and gender were not significantly different between controls and mild TBI.

### 3.2 Myelin water fraction (MWF)

Figure 1, meant for illustration purposes only, shows the MWF maps of six participants with TBI and six controls who were selected because they were closest in demographics. Note the overall qualitative reduction in MWF in each TBI participant. This was confirmed by a significant decrease in global white matter (GWM) MWF the mTBI group compared to controls (see first row of Table 2 and Figure 2). Analysis of the regions of interest (ROIs) for each group showed a significant decrease in MWF in the mTBI group compared with the controls in the majority of ROIs (see Figure 2 and Table 2), excluding the left anterior thalamic radiation (ATR) and right inferior longitudinal fasciculus (ILF) which were trending. After correcting for multiple comparisons using Bonferroni-Holm, only the splenium of corpus callosum (SCC) and left ILF were still significant. Due to the low subject numbers in the sTBI group it was not statistically compared.

**Table 2:** Mean and standard error of myelin water fraction (MWF) of all regions of interest; global white matter (GWM), splenium of corpus callosum (SCC), body of corpus callosum (BCC), genu of corpus callosum (GCC), anterior thalamic radiation (ATR), minor forceps (MN), inferior and superior longitudinal fasciculus (ILF and SLF respectively), corticospinal tract (CST) for controls and mTBI compared with an independent sample’s t-test. P-values below 0.05 are bolded and those that survive Holm-Bonferroni analysis for multiple comparisons are starred.

| Region of Interest |  | Control (n=12) | Mild TBI (n=18) | p-value |
| --- | --- | --- | --- | --- |
| GWM |  | 0.114±0.005 | 0.101±0.003 | **0.02** |
| SCC |  | 0.147±0.006 | 0.125±0.004 | **0.004\*** |
| BCC |  | 0.111±0.006 | 0.095±0.004 | **0.02** |
| GCC |  | 0.103±0.005 | 0.086±0.004 | **0.01** |
| MN |  | 0.087±0.006 | 0.074±0.002 | **0.02** |
| ATR | Left | 0.122±0.006 | 0.109±0.004 | 0.05 |
|  | Right | 0.112±0.006 | 0.097±0.003 | **0.03** |
| ILF | Left | 0.100±0.005 | 0.079±0.004 | **0.003\*** |
|  | Right | 0.133±0.006 | 0.120±0.004 | 0.05 |
| SLF | Left | 0.131±0.005 | 0.115±0.004 | **0.02** |
|  | Right | 0.144±0.006 | 0.126±0.003 | **0.01** |
| CST | Left | 0.205±0.006 | 0.189±0.004 | **0.03** |
|  | Right | 0.194±0.006 | 0.178±0.004 | **0.03** |

**Figure 1:**
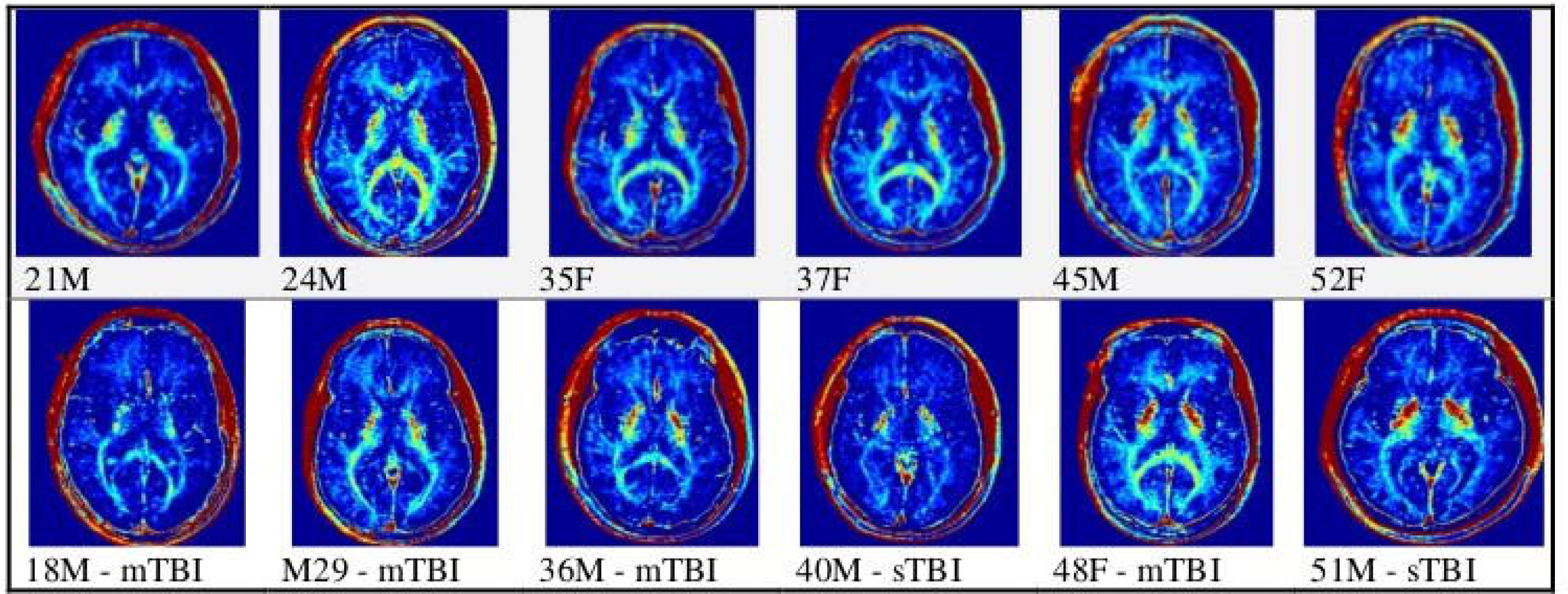
Sample myelin water fraction (MWF) maps for control and TBI participants.

**Figure 2:**
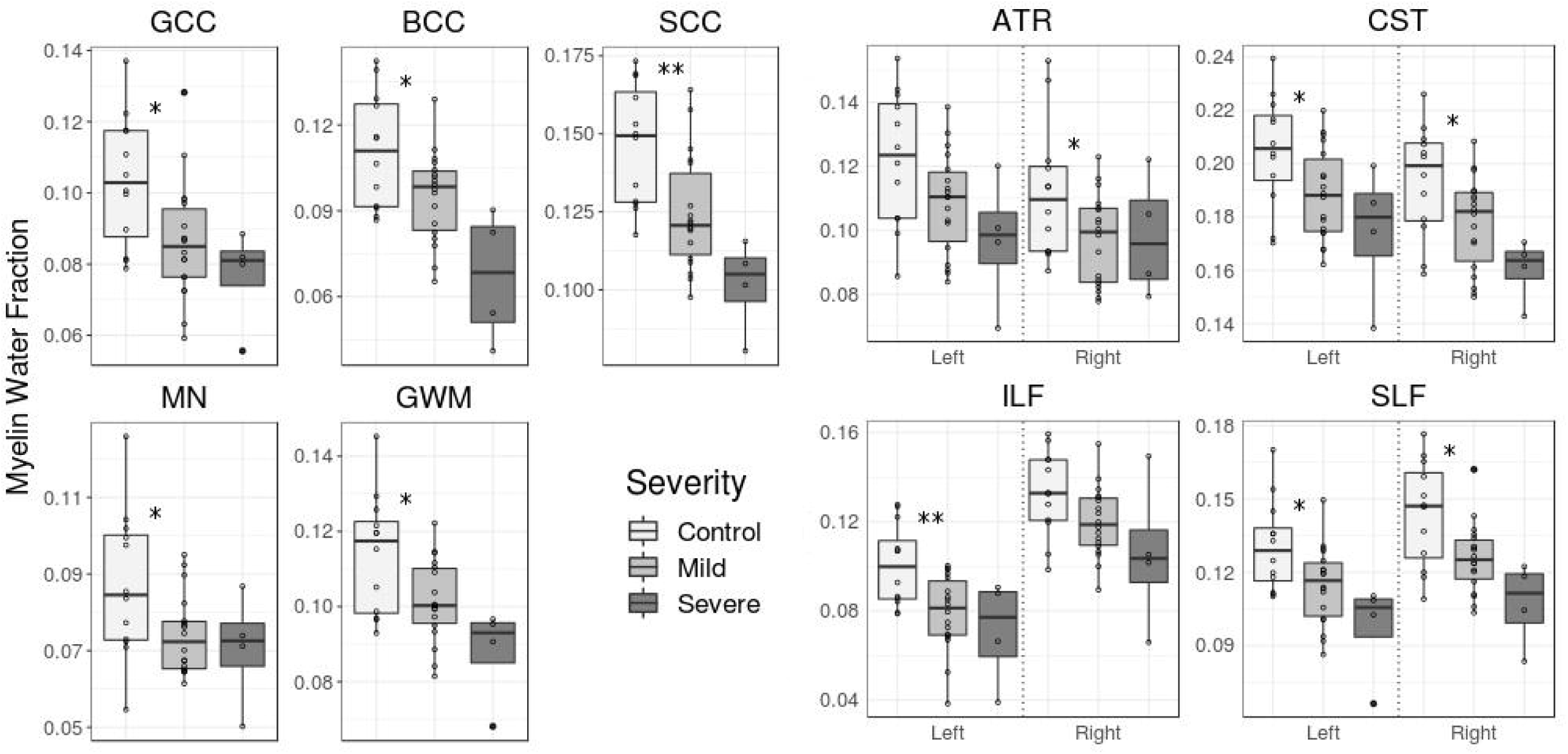
Myelin water fraction (MWF) for all controls and TBI subjects; in all regions of interest; genu of corpus callosum (GCC), body of corpus callosum (BCC), splenium of corpus callosum (SCC), minor forceps (MN), global white matter (GWM), left and right anterior thalamic radiation (ATR), superior and inferior longitudinal fasciculus (SLF and ILF respectively) and corticospinal tract (CST). Independent t-test comparison between control and mTBI uncorrected p-values are represented as *<0.05, **<0.01 and are taken from Table 2.

The relationship between age and years of education and MWF across all controls and mTBI were evaluated using Pearson’s correlations. Years of education was significantly positively correlated with MWF in all ROIs except in the SCC and left ILF and age was not significant in any ROI. Figure 3 shows MWF of global white matter in controls and mTBI plotted against years of education and age for each subject group. In the ANCOVA, group differences indicating lower GWM MWF in mTBI patients relative to controls were significant **p=0.04** after removing variances accounted for by years of education, which had a significant effect on the model (**p=0.01**), and age, which did not (p=0.4). After controlling for education and age, the following ROIs still showed significant group differences: GCC (**p=0.03)**, SCC (**p=0.01**), SLFL (**p=0.04**), and SLFR (**p=0.02**), the other ROIs were not significant, p>0.05.

**Figure 3:**
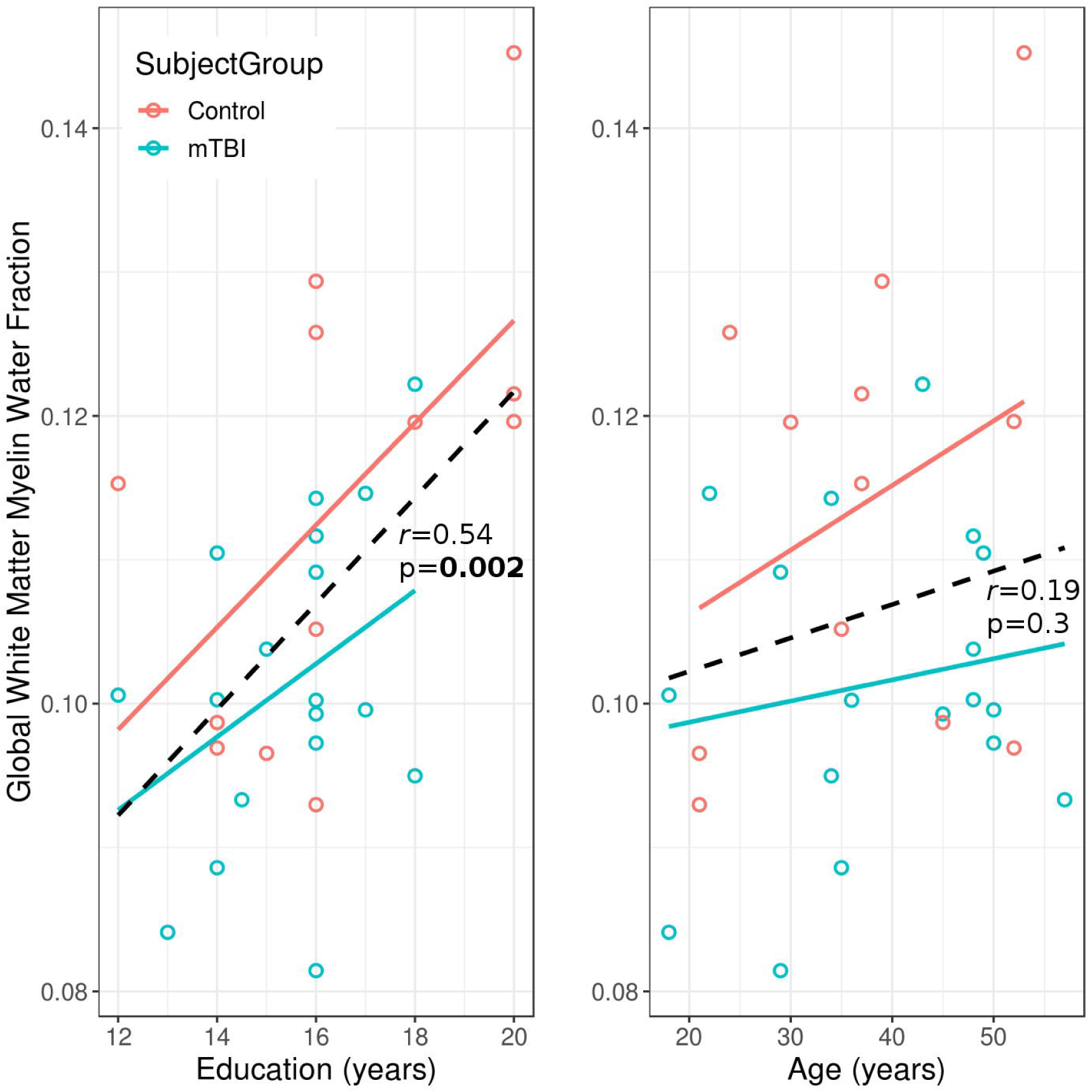
Effect of years of education and age on myelin water fraction (MWF) of global white matter across controls and mild TBI. The linear fit over controls and mTBI is represented by the black dashed line and the Pearson’s correlation coefficient (*r*) and p-value between education/age and MWF are reported.

### 3.3 Correlations between MWF and cognitive scores

Average age adjusted cognitive composite scores for Crystallized and Fluid cognition are given in Table 3 and the distribution of scores are given in Figure 4 for each subject group. Fluid cognition was significantly higher in controls than mild TBI, and both means were above 100. Crystallized cognition was not significantly different between groups and both means were above 115. The Cohen’s d suggested a moderate effect size for Crystallized and a large effect size for Fluid comparisons. Figure 5 shows the correlations between cognitive scores and GWM MWF for mTBI subjects. No significant correlations were found between mTBI MWF and cognitive scores in any ROI.

**Table 3:** Mean and sd of age-adjusted cognitive scores for controls and TBI with MRIs compared using an independent t-test and a Cohen’s d to measure effect size.

| Subject Group | N | Crystallized Cognition Composite Score |  |  | Fluid Cognition Composite Score |  |  |
| --- | --- | --- | --- | --- | --- | --- | --- |
|  |  | Mean (sd) | p-value | Cohen’s d | Mean (sd) | p-value | Cohen’s d |
| Control | 9 | 123(5) | 0.09 | 0.52 | 117(12) | **0.01** | 1.1 |
| mTBI | 18 | 116(11) |  |  | 101(17)<br>n=17 |  |  |
| sTBI | 4 | 107(11) |  |  | 85(6) |  |  |

**Figure 4:**
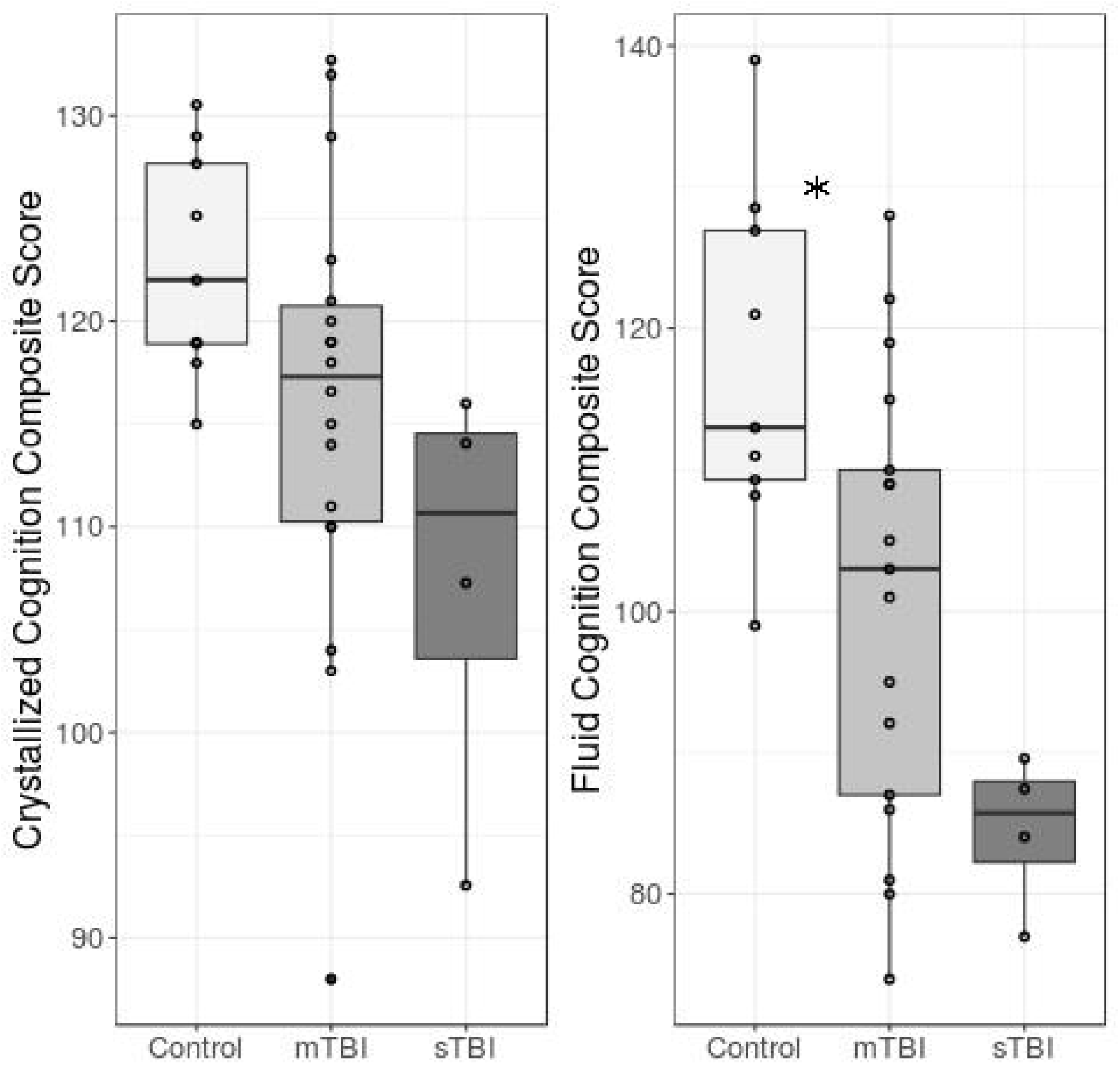
Age adjusted Fluid and Crystallized Composite scores for each subject group and subject, and independent t-test comparison between control and mTBI p-values are represented as *<0.05.

**Figure 5:**
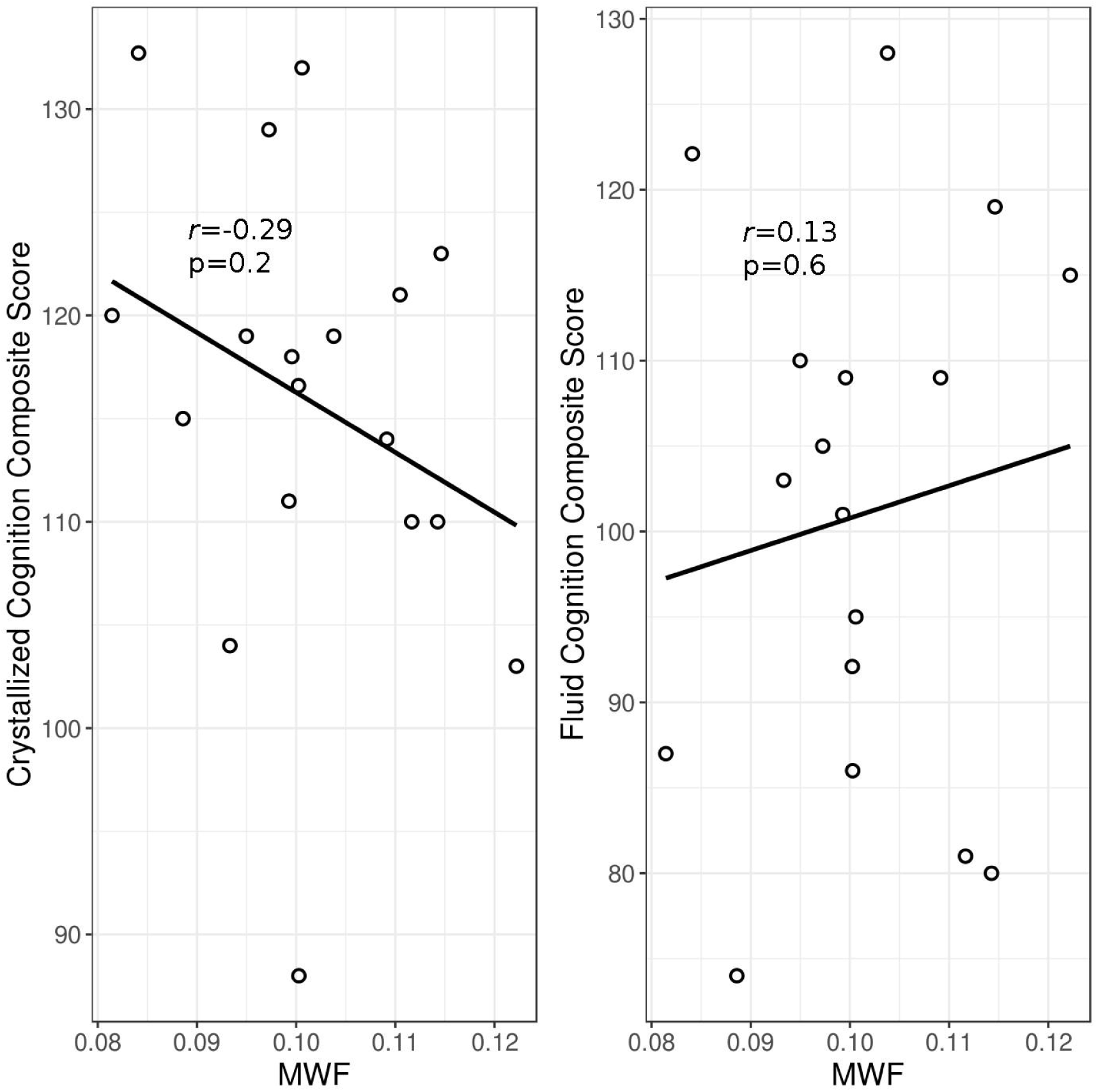
Age adjusted Fluid and Crystallized Composite scores compared to global white matter myelin water fraction (MWF) for mild TBI participants. Pearson’s correlation coefficient (*r*) and p-value between are reported.

## 4 Discussion

In this study, we report several new findings in relation to MWF in chronic TBI. First, we found significantly lower MWF in chronic mild TBI as compared to matched age, gender and education healthy controls. Lower MWF was evident globally, and in most white matter regions examined, however only two ROIs survived analysis for multiple comparisons, SCC and left ILF. For global white matter the difference in MWF between mTBI and control was 12% - for the corpus callosum this decrease was 15-18%. The consistent decrease in MWF in the participants with mTBI compared to controls in all areas examined is a strong result. The number of years of education had a significant effect on most ROI MWF measures but age did not. After accounting for the variance in MWF associated with differences in years of education, GCC, left ILF, SCC, SLF and global white matter MWF still exhibited significant group differences between controls and mTBI, indicating that the reduced MWF in patients was not solely due to education effects.

Severe TBI (sTBI) MWFs were generally lower than controls and mild TBI, however due to the low number of subjects in this category we did not able to statistically compare this group to the others. We would expect differences between controls and sTBI, as using a different technique to measure MWF, Choi et al.(70) also found significantly decreased MWF in a group with sTBI compared to healthy controls (these patients were only 2-months post-injury). However, the finding in mTBI is significant as most mTBI subjects are expected to return to pre-injury levels in terms of cognition and myelin in the chronic stage. For global white matter the difference in MWF between controls and mTBI was 12% - for the corpus callosum this decrease was 15.5-18%. A previous MWF study(53) on athletes with concussion found a decrease in SCC MWF in the acute stage but return to baseline MWF at a month 2 follow up. We found the mTBI group was significantly lower than controls in global white matter, all three regions of the corpus callosum, MN, right ATR, left ILF, SLF and CST. There is previous evidence that a subset of people with mTBI report persistent symptoms and problems. It is likely that our group of participants highlights those with ongoing issues(64–69), as most participants in our study self-reported symptoms consistent with post-concussive syndrome. This is further supported by the fact that we found significant cognitive differences between controls and mTBI.

The corpus callosum connects the two hemispheres of the brain and thus is prone to the twisting and sheering motion during a TBI. The corpus callosum, and specifically the splenium of the corpus callosum is highlighted in TBI studies as a main area of change and injury (86). As well, the splenium is an area known to be rich in myelin and thus a high MWF (30). We found the CC, and specifically the SCC (significant after correction for multiple comparisons) sensitive to detecting differences between controls and mTBI.

The exact pathway of myelin changes following TBI is not well characterized in humans, however, one post mortem human brain study found myelin loss (24). Many animal models have been developed to examine TBI (87–89). Disruptions of myelin sheaths have been found in stretching injury models of optic nerve in guinea pig (90,91). Recent studies in rodent models have shown extensive demyelination (22,23), remyelination but also, interestingly, extra myelin sheaths similar to redundant myelin which occurs during myelin development (92,93). This pattern of demyelination, and excessive myelin may make it difficult to separate controls from TBI in early phases of TBI, as one would decrease and the other increase MWF. There is some evidence that clearance of myelin is slow and may further hinder axon/myelin repair(94,95). However, there is also evidence of remyelination which would recover axonal signaling and functioning. Grin et al (25) reported that rats showed improved behavioral function following remyelination. Demyelinated axons are more vulnerable to damage, so ongoing loss of myelin can lead to further axonal damage, showing the importance of monitoring ongoing myelin changes post initial injury.

MWF has been found to change with age and years of education in earlier studies. Our study confirmed the relationship between MWF and years of education in healthy adults, however we also found this relationship in our TBI cohort. We did not find a significant correlation between age and MWF in either cohort or combined, which is different than what was found in the schizophrenia studies (56,58). This is most likely due to the much lower mean age in the schizophrenia study. MWF changes are more dramatic at lower ages, while the increase of MWF with age is gradual in middle age (57), and thus the influence of age may not have been detectable in our cohort. These findings underline the importance of matching groups not only on age and gender but also years of education.

TBI subjects showed lower Crystallized and in particular Fluid cognitive scores than controls, suggesting decreased cognitive functioning in patients relative to controls. Despite this, it should be noted that the mean cognitive scores were higher than or equal to 100 (average range) for both groups, indicating that in this study we have a highly educated group of patients and controls. We did not find a significant correlation between measures of MWF and either crystallized or fluid cognition for mTBI. A recent study reported a significant association between global MWF and processing speed in a sample of patients with moderate to severe TBI at 3 months post-injury(70). We did not have a sufficient sample size in sTBI to evaluate relationships between cognitive scores and MWF. Persistent cognitive deficits are rarely found in mTBI; however, here we found lower cognitive scores in mTBI than controls, and this may be due to the select group of mTBI we studied who have persistent self-reported symptoms. Thus, the findings of this study may not be generalizable to patients with mild TBI who do not present with persisting symptoms.

Our TBI sample was highly educated and their mean CC score was above average which, may signal an elevated level of cognitive reserve. This high cognitive reserve may, limit the potential amount of cognitive morbidity, i.e. these patients may be able to tolerate more injury to the brain without it affecting their cognition. This reserve may prevent or restrict the observation of cognition-MWF associations(96). These associations would be predicted to be more evident in patients who were less educated or had less cognitive reserve.

Traumatic brain injury imparts a huge burden on society and even mild TBI can have long-term effects on patients. There is increasing evidence that TBI is not a ‘single event’ but rather an ongoing process which unfolds across months, years, and possibly over a lifetime; thus, improved monitoring of long-term effects of TBI is essential (97). White matter tracts are particularly vulnerable to a TBI and monitoring the myelination of these tracts may expand our knowledge and inform the development of diagnostic and prognostic biomarkers. Myelin water fraction is a powerful quantitative technique to examine myelin content and has been used to examine a variety of brain diseases and disorders (41). Here we show MWF differences in mTBI compared to healthy controls, months to decades post-injury, and show the utility of MWF to examine long-term effects of mild TBI.

## 5 Acknowledgements

This project is funded by MITACS in partnership with the Eaton-Arrowsmith group.

## 6 Author Contributions

I.T., W.P., A.L.M. and N.V.B. planned the study. B.R-S completed data analysis and prepared the manuscript. A.M.M. V.P. and L.B. managed the data collection. All authors contributed to the interpretation of the results and reviewed the manuscript.

## 7 Conflict of Interest Statement

Dr. Torres has served as consultant for Lundbeck Canada and Sumitomo Dainippon.

Dr. Panenka has a forensic psychiatry practice that focuses on head trauma.

Dr. Zhang is a GE clinical scientist but was not working with GE at the time she completed the research for this paper.

Dr. Virji-Babul: “No competing financial interests exist”

Dr. MacKay: “No competing financial interests exist”

Dr. Vavasour: “No competing financial interests exist”

Ms. Russell-Schulz: “No competing financial interests exist”

## 9 Data availability statement

The UBC Ethics Board will not allow us to store data outside of Canada, however if there are researchers interested in using the data we can apply for an amendment to share anonymized data with individual institutions.

Information on implementing the myelin water imaging technique and analysis code is available on github via a request form through the UBC MRI Research Centre website (https://mriresearch.med.ubc.ca/news-projects/myelin-water-fraction/).

