## Supplemental Table 1a for "Myelin water fraction decrease in mild traumatic brain injury"

Supplementary Material

Table 1b: Traumatic brain injury (TBI) subject injury description and demographics

| Subject | Approx. Time Since Injury (years) | Age (years) | Gender | Years of Education | Injury Type | Injury Severity |
| --- | --- | --- | --- | --- | --- | --- |
| S1 | 21 | 51 | M | 14 | MVA | Severe |
| S2 | 28 | 54 | F | 18 | MVA | Severe |
| S3 | 23 | 36 | M | 16 | Multiple Concussions | Mild |
| S4 | 7 | 40 | M | 12 | MVA | Severe |
| S5 | 1 | 18 | M | 13 | Multiple Concussions | Mild |
| S6 | 7 | 50 | F | 16 | MVA | Mild |
| S7 | 7 Months | 29 | M | 16 | Multiple Concussions | Mild |
| S8 | 2 | 43 | F | 18 | MVA | Mild |
| S9 | 2 | 45 | M | 16 | Multiple Concussions | Mild |
| S10 | 2.5 | 48 | F | 15 | Assault | Mild |
| S11 | 1.5 | 57 | F | 14.5 | Fall | Mild |
| S12 | 4 | 34 | F | 18 | MVA | Mild |
| S13 | 1 | 22 | M | 17 | Fall | Mild |
| S14 | 23 | 45 | M | 14 | Blast | Severe |
| S15 | 1 | 34 | F | 16 | Sport | Mild |
| S16 | 6 months | 29 | M | 16 | Sport | Mild |
| S17 | 2 | 35 | M | 14 | Assault | Mild |
| S18 | 1 | 50 | M | 17 | Fall | Mild |
| S19 | 2 | 18 | F | 12 | Sport | Mild |
| S20 | 21 | 49 | F | 14 | MVA with brain infection | Mild |
| S21 | 9 months | 48 | M | 16 | Sport | Mild |
| S22 | 5 | 48 | F | 14 | Multiple Concussions | Mild |

MVA – motor vehicle accident (single incident)
